# Optimized Mappings from Biological Hip Moment Estimates to Exoskeleton Torque can Personalize Assistance Across Users and Generalize Across Tasks

**DOI:** 10.1101/2025.08.29.671780

**Authors:** Justine C. Powell, Ethan B. Schonhaut, Dean D. Molinaro, Aaron J. Young

## Abstract

Recent advancements in data-driven methods have enabled real-time estimation of biomechanical states for exoskeleton control. While biological joint moments can be directly used to scale exoskeleton assistance, this approach is often suboptimal. An optimized mapping between biological joint moments and exoskeleton assistance could enhance end-to-end controllers based on the user’s physiological state. We introduce a flexible parametrization of biological moment-based control using delay, scaling, and shaping terms to transform joint moment estimates into commanded torque. We performed human-in-the-loop optimization, using metabolic cost to evaluate each iteration’s controller parameters, for 9 subjects across three ambulation modes: level walking at 1.1 m/s, 1.5 m/s, and 5° inclined walking. We evaluated three methods of exoskeleton control: 1. Personalized/Task Dependent, 2. Task Dependent/Non-personalized, and 3. Task Agnostic/Non-personalized. On average, our personalized approach provided the greatest benefit of 18.3% reduction in metabolic cost compared to walking without the exoskeleton, with the task dependent and task agnostic controllers producing similar reductions of 8.6% and 8.4%, respectively. Our results show that while generalizable, task agnostic control parameters can improve user energetics across cyclic tasks, fully personalized exoskeleton control parameters yield larger metabolic reductions, highlighting the value of personalizing exoskeleton assistance to users across many diverse tasks.

## I. INTRODUCTION

Artificially intelligent technologies capable of improving everyday life are becoming increasingly commonplace in society. Wearable devices, such as robotic assistive exoskeletons, can augment user movement to reduce the amount of energy expended throughout the day. These devices, however, are difficult to deploy outside of the lab due to human and task level variability, prompting the need to personalize exoskeleton assistance to the user. Humans continually adapt and optimize our movements to successfully improve task efficiency, as evidenced during steady state treadmill walking [1]. Even with a reduced task space that only encompasses ambulatory tasks, the conditions users may adjust to throughout the day are numerous, with every individual performing these adaptations in unique ways. Assistive robotic exoskeletons have been shown to reduce the energetic demands of human locomotion across discrete ambulatory tasks [2], [3], [4], however these studies often do not scale to the wide array of movements or environments users may encounter throughout the day. For wearable assistive exoskeletons to provide maximum benefit, they must allow users to fine tune and personalize these devices to fit their needs, while simultaneously allowing for adaptability and generalizability to the vastly diverse nature of human movement.

Key barriers to translating wearable exoskeletons from the lab to the real world lie in developing control strategies able to handle the transient nature of human mobility. Current exoskeleton control strategies can generally be broken up into three hierarchical levels: high, mid, and low level control [5], [6]. High level control is used to detect and characterize the environmental conditions the user is operating in, such as walking speed or ground slope or the internal state of the user, such as biological joint moment. Mid-level controllers, such as the control framework described in this manuscript (Fig. 1a), typically use environmental information from a high-level controller in conjunction with real-time human-based sensor data to compute the amount of exoskeleton joint torque based on a set of state dependent dynamics. Low level controllers are device dependent [6] attempt to minimize the error between the commanded torque from the controller and the actual torque delivered by the actuators to the subject during deployment.

**Fig. 1:**
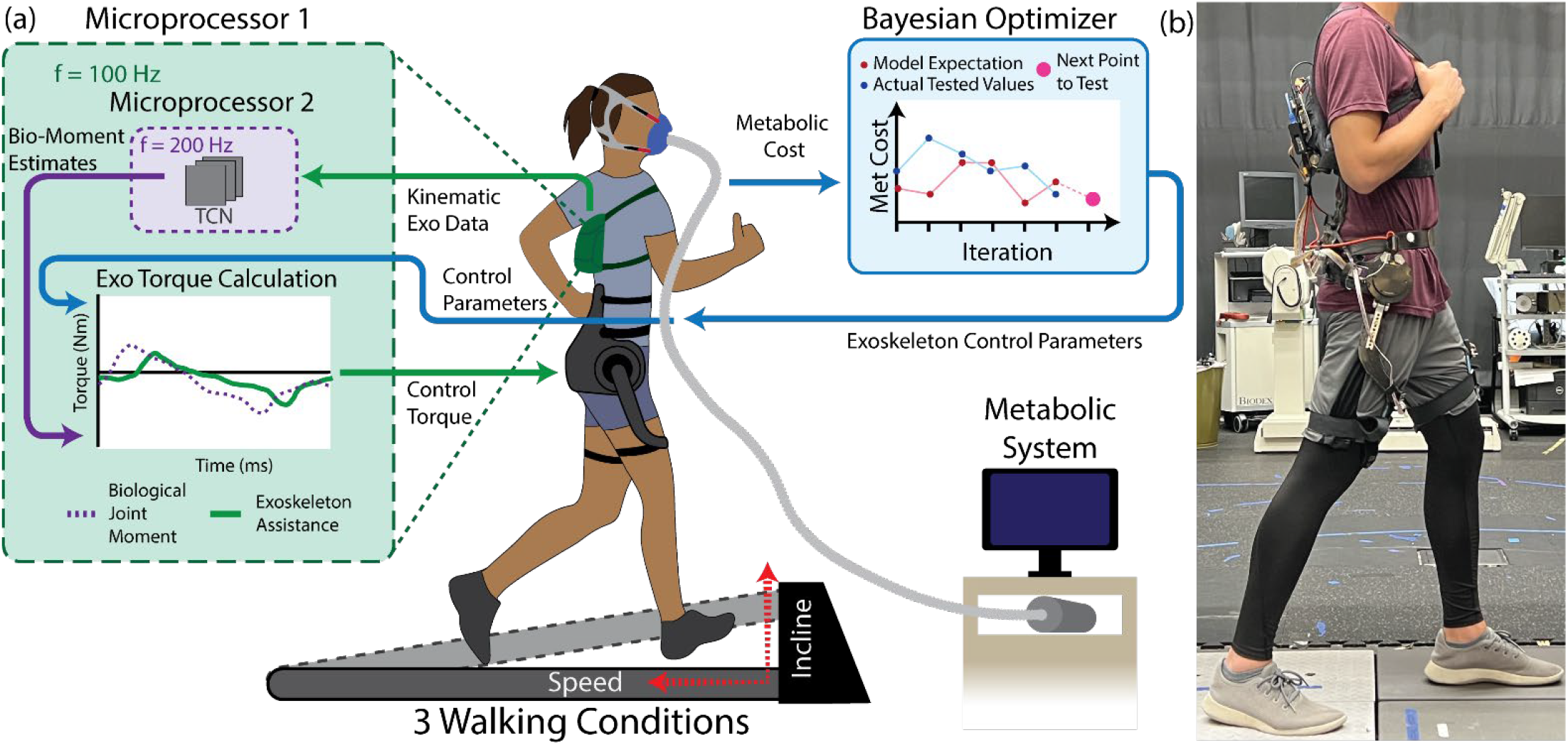
**(a)** Schematic of the experimental design. The experiment was run across three different walking conditions: level ground walking at 1.1 m/s, level ground walking at 1.5 m/s, and 5° incline at 1.1 m/s. The human in the loop optimization (HILO) loop, in blue, was used to metabolically optimize the exoskeleton’s control parameters over the course of 21 iterations. Two microprocessors (green and purple, respectively) are used to control the exoskeleton’s actuators. A National Instruments MyRio (Microprocessor 1) receives information directly from the exoskeleton’s actuators and peripheral sensors and sends the information to an Nvidia Jetson Nano (Microprocessor 2). The Jetson Nano feeds the information through a temporal convolutional network (TCN) to estimate the subject’s biological hip joint moment and returns the information to the MyRio. Biological joint moment estimates and the control parameters from the HILO loop are then passed to the MyRio to calculate the instantaneous torque command to send to the actuators. **(b)** Side view of the hip exoskeleton used during experimentation consisting of a custom carbon fiber hip interface, colocated hip motors, and an electronics backpack containing the system mechatronics. Note, the subject in (b) is an author on this manuscript and has thus consented to the use of this image.

Mid-level lower limb exoskeleton controllers are diverse in their formulation and are characterized by the modality of their input [5], [6]. Kinematic based controllers transform the subject’s kinematic information, measured via sensors such as encoders or inertial measurement units (IMUs) placed on board the exoskeleton, to exoskeleton torque using an equation such as impedance control, the user’s gait cycle, or a pre-prescribed spline-based timing controller [3], [7], [8], [9]. These controllers, however, are highly task specific and are only successful under strict conditions, which does not accurately represent the diversity of movements and environments that humans operate within. Neural based controllers use sensors that capture signals inside of the human subject to compute exoskeleton assistance. This includes controllers that transform muscle activation measured via electromyography (EMG) to output torque [10], [11], as well as more invasive techniques that transform brain activation via electroencephalography (EEG) to exoskeleton assistance torque through the use of brain computer interfaces [5], [6], [12]. While these approaches directly use the user’s physiological signals to influence assistive device control, they can be difficult to incorporate in real-time due to issues with system reliability and robustness [13], [14], and often still require task specific gains and thresholds. A relatively newer class of control uses deep learning (DL) to transform data from the subject sensor domain to the actuator torque domain, usually through a proxy that estimates some internal physiological state of the user, such as biological joint moment [15], [16], [17].

Computing internal physiological states such as lower limb biological joint moments, however, is computationally expensive, often requiring access to motion capture systems and ground reaction forces to compute ground truth biological moment labels [18], [19]. Likewise, assistive exoskeletons often rely on onboard sensing, making real-time computation of these labels within the control loop of a device extremely difficult. Deep learning-based estimators can circumvent this computational limitation by estimating biological joint moments instantaneously from sensors that are easily deployed onboard a wearable assistive device. These approaches, such as the temporal convolutional network (TCN) developed by Molinaro *et al.* [15], [16] have been shown to be an effective method of continuous exoskeleton control, or end-to-end control, across a wide range of users and tasks [15], [16], [17], [20]. This approach can directly estimate biological joint moments at the hip and knee based on different wearable sensors such as joint encoders, IMUs, and pressure sensing insoles. Furthermore, this approach uses a task and subject agnostic approach by training on a sufficient number of diverse subjects and tasks to improve generalizability across different users and activities [15], [16]. This study employs the TCN developed by Molinaro *et al.* [16], where biological joint moment estimates are produced instantaneously upon receiving exoskeleton sensor data. The estimates from this network are inputs into a mid-level controller that scales, delays, and filters torque estimates to compute the assistance applied to the user based on the user’s body mass and the maximum torque output of the device actuators. The availability of this instantaneous sensor data along with new developments in mid-level control parametrization provide an avenue to use biological torque estimates as a control signal for optimized and improved exoskeleton assistance.

While estimating biological joint moment represents an effective method of generalizable exoskeleton control across different tasks and activities [15], [16], studies have shown that providing purely biological joint moment-based assistance is not metabolically optimal during steady state walking [8], [9]. Similarly, pilot studies from our previous work have shown that delaying exoskeleton assistance relative to the biological hip moment improved metabolic benefit across subjects [16], [20]. During these pilot studies, this benefit was expected because the added delay maximizes the positive work done by the exoskeleton [21], [22], indicating that further optimized parametrizations of exoskeleton mid-level controllers can increasingly improve metabolic benefits for users. Human in the loop optimization (HILO) represents a promising approach to determine the best controller settings and parameters on a per user and per task basis in steady state, cyclic tasks such as walking. This process uses a human outcome measure or cost, such as user energy expenditure, as a metric to assess the effectiveness of a given controller’s settings or parameters. These parameters and their associated cost are then fed into an optimizer that determines the next set of parameters to test [8], [9], [21], [22], [23], [24]. As the optimizer iterates, the controller parameters that correspond to the minimum cost for the user and task can be found. Most HILO torque assistance profiles, however, are based on prescribed spline-based torque profiles for a specific task, which does not account for the diversity of human movement. These devices often require a new optimization for each task-specific assistance profile provided by the device, which is not only time consuming to perform, but most importantly ignores the link between the device and the user’s underlying physiology.

This study expands on previous work from our group by Molinaro *et al.* [16] by formally optimizing the relationship between biological joint moment and applied exoskeleton torque across multiple subjects and ambulation modes (Fig. 1). Specifically, we aim to understand how sensitive exoskeleton mid-level control is to different parametrizations of the applied continuous torque assistance of our personalized approach during steady state locomotion tasks. Furthermore, we aim to analyze the relationship between positive exoskeleton power and metabolic cost for our continuous end-to-end control approach, as well as how user preference responds and changes with different exoskeleton control parameters. We hypothesize that **I:** Personalized, task specific biological-torque based exoskeleton control will provide a significant energetic benefit in comparison to non-personalized, task dependent controller settings, **II:** non-personalized, task dependent controller settings will provide significant metabolic benefit over non-personalized, task-agnostic controller settings, **III:** metabolic cost reductions will be correlated with positive power provided by the exoskeleton during assistance, such that the most effective strategies maximize positive mechanical work provided by the exoskeleton to the user. Here, we introduce our control framework that leverages the advantages of a generalizable, task-agnostic end-to-end exoskeleton controller in conjunction with the benefits of HILO to optimize and unify the user’s underlying physiological signals to end-to-end exoskeleton control.

## II. Methods

### A. The Exoskeleton & Controller

The device used in this experiment was a fully autonomous, one degree-of-freedom (DOF), untethered hip exoskeleton (Fig. 1b) that was designed, fabricated, and previously validated by our group [16]. The exoskeleton’s assistance was designed to assist in the sagittal plane (hip flexion and extension), with torque assistance supplied by actuators with a max output torque of 18 Nm and transmitted to the subject’s lower body through custom carbon fiber thigh struts. The exoskeleton’s onboard sensors included IMUs placed posteriorly on the pelvis and on the frontal plane of each thigh, as well as encoders available via the device actuators. The backpack attached to the exoskeleton contained batteries, two microcontrollers, and electrical breakout boards required to control the exoskeleton. The total weight of the system was 4.8 kg, with more details on device design available in our previous publication [16].

This study’s goal was to determine the optimal relationship between biological joint moment and output exoskeleton torque at the hip joint. Previous work by our group has validated the use of a subject independent TCN model to directly estimate biological joint moments as a means of exoskeleton control using the same hip exoskeleton used in this study with an R^2^ of 0.840 ± 0.045 and an RMSE of 0.142 ± 0.021 N·m/kg) [16]. This TCN model was deployed on an Nvidia Jetson (Nvidia, Santa Clara CA) microprocessor which communicates with an NI MyRio (National Instruments, Austin TX) during the experimental loop (Fig. 1a).

### B. Controller Parametrization

The mid-level control parametrization (1) uses three terms to transform estimated biological hip moment 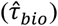 to exoskeleton assistance torque (*τ_cmd_*) at a given time (t). The scale (*α*) term changes the magnitude of torque assistance as the actuator’s maximum torque is much less than that of the hip joint. In this experiment, the scale term was not optimized and instead held constant at 20% of the subject’s biological hip joint moment, due to device torque limits. The delay term (*d*) determines the time difference in milliseconds between when the subject exerts a biological joint moment and when they receive the exoskeleton assistance. In Molinaro *et al.* [16], the delay term was held to 125 ms based on an N=3 pilot study comparing metabolic cost to several delay terms, which furthermore, was in correspondence to the hypothesis proposed in Camargo *et al.* [23] that timings between 100 to 150 ms could maximize work the exoskeleton supplies to the user. The shape term (λ) is a non-linear transformation based on a power law to change the impulsiveness of the torque profile. The values of λ were chosen such that during the optimization process and within any ambulation mode, all combinations of shape and

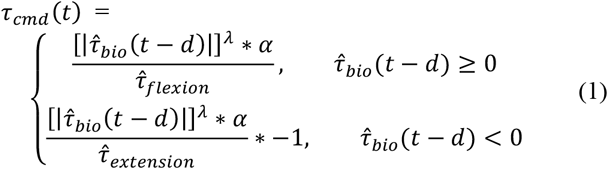

delay parameters would result in similar magnitudes of *τ_cmd_*. However, because the shaping term is exponential, with values ranging from 0.5 to 1.6, there are instances where the magnitude of commanded exoskeleton assistance torque exceeds the capability of the device actuators. To account for this, we estimated the peak net, flexion, and extension joint moments for each subject during a habituation period and adjusted the scaling term, α, to ensure a consistent scaling of 20% peak biological joint moment of the user as parameters are optimized during the experiment. Furthermore, due to device limitations, any torque commanded above peak exoskeleton actuator torque was limited to 18 Nm of torque applied to the user. The effects of each term on an optimal biological joint moment profile adapted from [9] can be seen in Fig. 2.

**Fig. 2:**
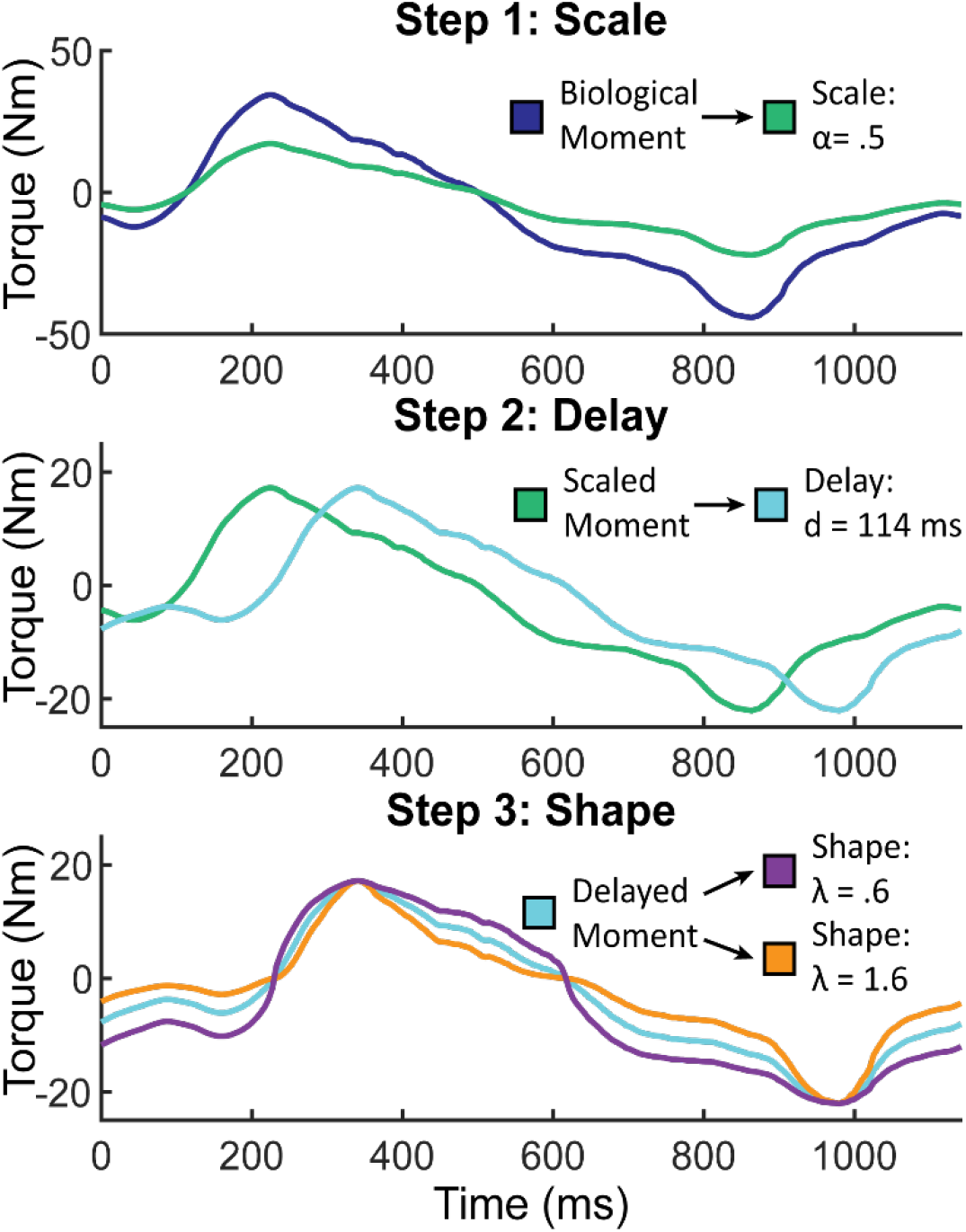
Steps to transform a subject’s biological hip joint moment to exoskeleton assistance torque. The subject produces the biological torque which is measured via onboard sensor data by the MyRio microprocessor. The Nvidia Jetson receives sensor data from the MyRio and passes it through a biological joint moment estimator where it is filtered and returned to the MyRio with a small filtering and communication delay. **Step 1:** A scaling factor *α* < 1 (α ≈ 0.2) reduces the magnitude of the moment to an actuator torque value within the system limits. **Step 2:** A delay term 65 *ms* ≤ *d* ≤ 240 *ms* (including the filter & communication delay) further shifts the exoskeleton control profile away from the original biological hip moment. **Step 3:** A shaping term. 5 ≤ λ ≤ 1.6 modulates the sharpness and flatness of the peaks of the control profile.

### C. Data Collection

Nine able bodied participants (8 male, 1 female, average mass 69.14 ± 13.83 kg) were recruited to participate in this study. The study experiment included ambulating while wearing the exoskeleton as the exoskeleton controller parameters were optimized across three different treadmill conditions in three separate 3-hour long sessions. The three treadmill conditions are as follows: level ground walking at 1.1 m/s, level ground walking at 1.5 m/s, and 5° incline at 1.1 m/s. Each session began with a 15-minute habituation period in which the subject was exposed to a range of parameter combinations to condition them to walking in the exoskeleton. The subjects then walked for 25 two-minute recorded trials consisting of two trials with no exoskeleton, two trials walking in the unpowered exoskeleton, and 21 trials walking with exoskeleton assistance. Prior to participating, each participant gave informed written consent to participate in a Georgia Tech Institutional Review Board-approved study.

To optimize our control framework, we used metabolic cost measured via indirect calorimetry as our cost function, which measures the user’s oxygen intake and carbon dioxide output to compute their energy expenditure [2], [25], [26]. Indirect calorimetry remains the gold standard for measuring user energy expenditure, making it an easily attainable metric to use during HILO. Our optimization occurred for each ambulation mode across 21 distinct trials, where the exoskeleton assisted the user. During each trial, we collected 90 seconds of sensor data from onboard the exoskeleton and two-minute measurements of metabolic cost via the Parvo metabolic cart (TrueOne 2400, ParvoMedics). Across all subjects and conditions, the first 6 iterations of assistance were made up of pre-selected parameters to seed the optimization process with the same data points across subjects (Fig. 3a). One of these 6 points, the purple triangle in Fig. 3, was selected based primarily on an N=3 study that swept only the delay term for biological joint moment control [16]. A second pre-determined parameter combination, represented by the orange triangle in Fig. 3, was chosen using an offline global optimization of the controller to another optimal control profile, as determined using spline-based exoskeleton controller [8], [9]. Real biological torque data along with a built-in global optimization function in MATLAB (The MathWorks Inc., Natick, MA) was used to determine the shape and delay terms from (1) that minimized the root mean squared error (RMSE) between the optimal control profile from Franks *et al*. [9] and a control profile generated using the parametrization from (1). The final four pre-selected parameters (pink, light blue, green, and dark blue triangles in Fig. 3) were chosen to provide a broad sampling of the parameter space to seed the Bayesian optimization process. In each session, the order of the 6 pre-selected points was randomized to mitigate ordering effects and reduce bias between subjects. Note that these 6 control parameters were used to habituate the subject to the exoskeleton and controller during the habituation session, as well as to determine the scaling term α used in the mid-level parametrization.

**Fig. 3:**
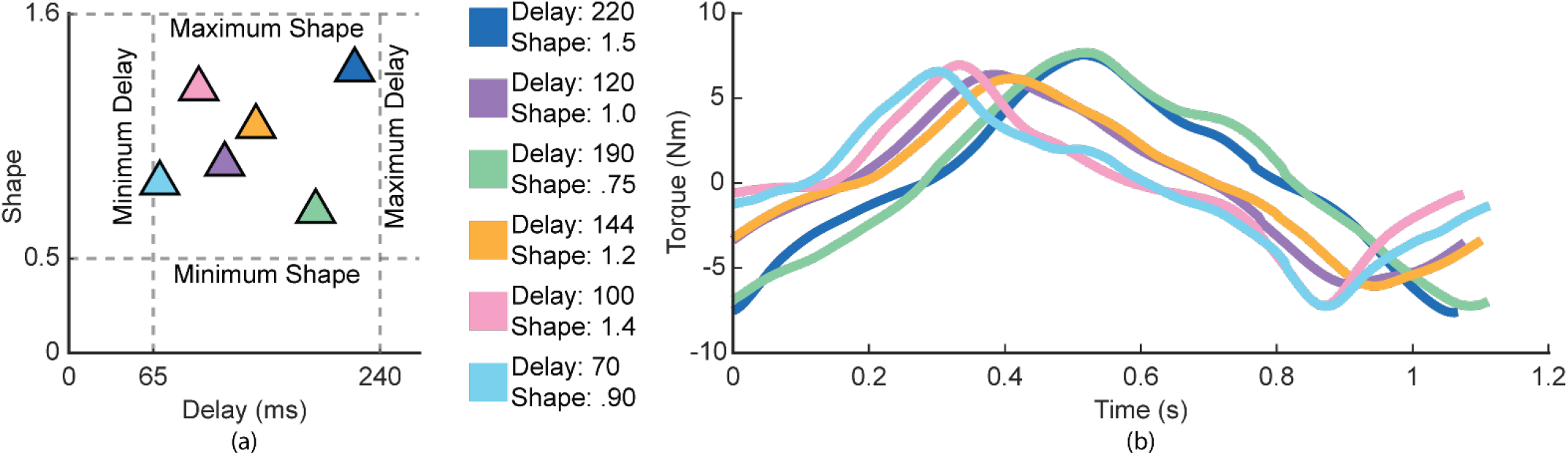
**(a)** Graphical representation of the exoskeleton control parameter space highlighting the two optimization parameters, shape and delay, for level ground walking at 1.1 m/s. The grey dotted lines represent the minimum and maximum values of each parameter that bounded the human-in-the-loop Bayesian optimization. These values were determined during an N=3 pilot testing of the experiment (apart from the minimum delay which is limited by the time it takes to estimate biological moments). The triangles represent the six control parameter combinations that were used to initiate the Bayesian optimization for each subject and every ambulation mode. The starting points were selected as they represent a wide range of possible control profiles, as well as 2 points that are near optimal points found in previous studies. **(b)** The exoskeleton assistance torque profiles that are generated by the control parameters in (a).

### D. Bayesian Optimization

We used Bayesian optimization to select the next fifteen iterations of assistance parameter combinations. Bayesian optimization was chosen as it has been found to be a time efficient method of HILO, and has outperformed previously used methods such as gradient descent [24]. Furthermore, Bayesian optimization suits HILO problems well as it is sample efficient and can directly be controlled to motivate exploration or exploitation to avoid falsely selecting local minima. When the exoskeleton control parameters were updated between iterations, the subject was allotted 30 seconds to adjust to the new parameters prior to beginning collecting data for the next optimizer iteration. Additionally, the subject’s preference between the current and previous iterations’ assistance was queried after each parameter change. Due to the metabolic mask required by the TrueOne 2400 Parvo system, the subject signaled better, worse, or no difference using hand signals during walking to avoid talking. Due to the physical demands of the data collection, subjects had predetermined scheduled break periods after every six iterations or by subject request and were allowed to rest until ready to continue.

Custom MATLAB code using the Gaussian process regressor (GPR) package was created to run the human-in-the-loop Bayesian optimization process during the experiment. The acquisition function used was an expected improvement equation with an additional restriction so that a parameter combination too close to any of the previously tested parameter combinations could not be selected [25]. We restricted the optimizer to selecting delay terms between 65 ms and 240 ms and shape terms between.5 and 1.6 (Fig. 3a); these bounds were determined to be within reasonable range of comfort in pilot testing of the controller. The optimization model was set to be moderately explorative rather than exploitative so that similar parameters weren’t tested repetitively and to improve model convergence within the 21 iterations (Fig. 4).

**Fig. 4:**
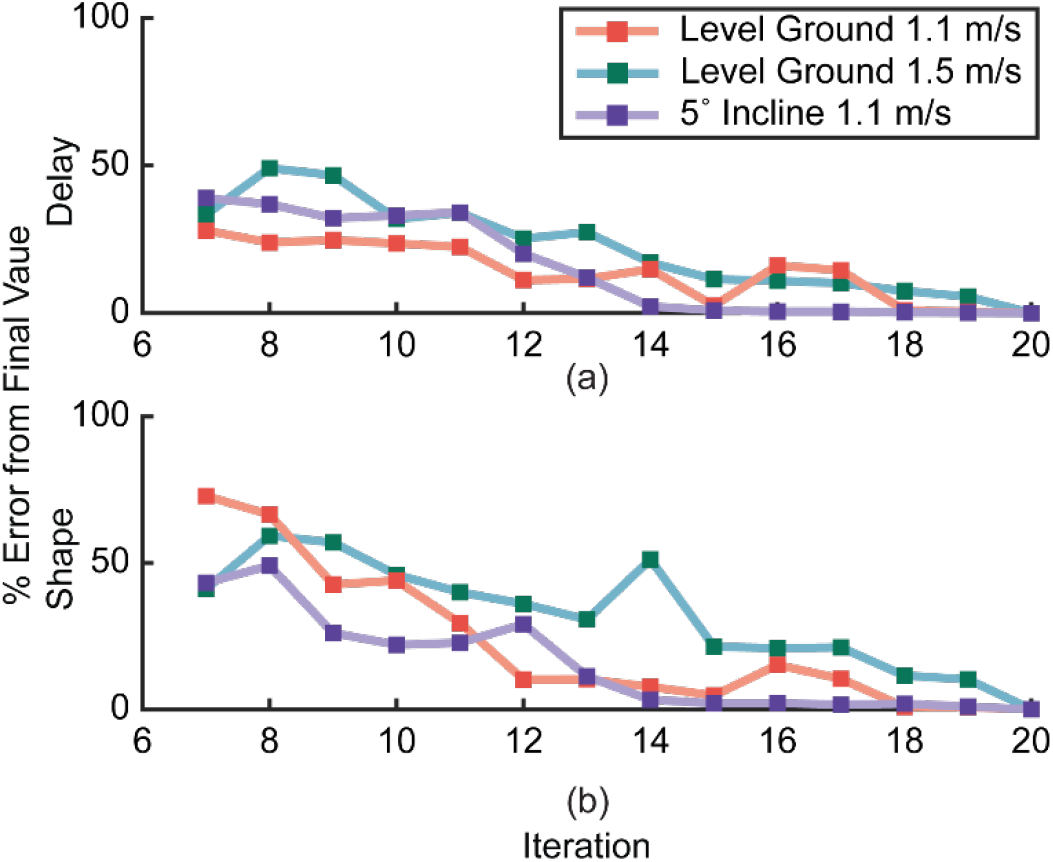
Convergence analysis of the human-in-the-loop Bayesian optimization across all three ambulation modes. Each plot begins at iteration seven after the six initial pre-selected parameter combinations. The model’s convergence to the final **(a)** delay and **(b)** shape values in absolute percent change between the Gaussian process regressor’s (GPR) believed optimal parameter and the actual value of the optimal parameter, i.e., the percent error at iteration 20 is equal to 0. For both the shape and delay parameters, the model was able to converge in fewer iterations during the inclined walking trials in comparison to the level ground trials.

### E. Metabolic Cost

Standard metabolic experiments typically require six minutes of data to compute metabolic cost [26], however due to the physical demand of our experiment, we collected two-minutes of metabolic data where the steady state metabolic cost value was estimated via a first order dynamic model applied to the two minutes of transient metabolic data, representing the current gold-standard within the field [27], [28]. This approach has previously been validated during metabolic and HILO studies that have shown that accurate estimates of metabolic cost require a minimum of one minute of respiratory data [26], [27]. To make comparisons across both subject and walking conditions, we normalized the metabolic data by the no exoskeleton metabolic value using (2), where all metabolic analyses are presented in units of percent change from the no exoskeleton condition (%Δ No-Exo).

For each subject and walking condition, we created metabolic landscapes using the controller parameters and their corresponding normalized metabolic cost from the 21 exoskeleton assisted iterations within each session using cubic interpolation and bounded radial basis function (RBF) extrapolation in areas that were outside of the convex hull of the data points tested. Each subject’s best parameters, as approximated by their specific metabolic landscape, are denoted on individual condition surfaces as well as the mean best parameters for that condition. The average condition surface shows the overall best parameter set across all tasks, as well as the best parameters for each task averaged across all subjects.

We used a one-way ANOVA in SPSS (IBM, SPSS Inc., Chicago, Illinois) to compare the metabolic cost across different best-case control parameters conditions: personalized control, task dependent control, and task agnostic control. We used the 27 metabolic landscapes determined from our experiment (9 subjects, 3 conditions) to find a single-best case parameter set, consisting of a shape and delay value, for each of the three conditions. Personalized control is both task and subject dependent, with best-case parameters corresponding to the lowest metabolic cost observed on the landscape for a specific subject during a specific task. Task dependent control is solely dependent on a specific task and is not personalized to the user, where parameters for a given subject are determined by averaging the metabolic landscapes from the 8 remaining subjects within that task. Task dependent control parameters are

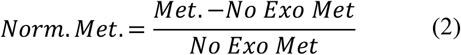

then derived from the corresponding lowest metabolic cost parameters on the task-specific subject-withheld landscape. Task agnostic control is independent of both subject and task, where a given subject’s metabolic landscapes for all tasks are withheld and the landscapes from all remaining subjects across all tasks are averaged into one surface.

Task agnostic control parameters are then derived from the corresponding lowest metabolic cost parameters found on the subject-withheld average surface. All percentage changes are with respect to the metabolic cost when the subject was not wearing an exoskeleton (No Exo), and all standard deviations are using a 95% confidence interval. Significant differences between both the no exoskeleton condition and between bars within an ambulatory condition cluster were determined using a one-way ANOVA with an alpha level of 0.5 and a multiple comparisons test with a post hoc Bonferroni correction. We compared between metabolic cost, torque, and power using linear regression, where we computed the slope, Pearson’s correlation coefficient (r), and p-values generated to determine the significance between each condition and trend.

### F. User Preference

We gave numeric values to the pairwise preference comparisons of the control parameters queried from the subjects during the optimization process. For a given chain of 21 control parameters and corresponding pairwise comparisons, every time a control parameter was rated as worse than another it lost one point of value and every time a control parameter was rated as better than another it gained one point of value. If a subject rated two parameter sets as equal, no points were either gained or lost for both parameter sets. Once every control parameter in each 21-iteration trial was given a numerical value, we created a preference landscape for each task and the overall task average, with a similar process of making the metabolic landscapes based on previously validated preference characterizations [29]. We then separated the parameter space into four quadrants to determine if controller preference could be reasonably predicted using this method. Across all modes, the 20 pairwise comparisons across 9 subjects were grouped by which two quadrants they were comparing, and the percentage of times the subjects preferred one quadrant to another was calculated. Furthermore, we computed Pearson’s correlation coefficient between the all-task average preference surface and the all-task average metabolic surface to evaluate if higher reductions in metabolic cost trended with control parameters that were more highly preferred by users.

### G. Exoskeleton Power & Torque

We analyzed how the power and torque provided by the exoskeleton trended with metabolic cost reduction. Average torque, peak positive power, and average positive power were calculated using unilateral data from the exoskeleton’s sensors. Torque (τ) is directly proportional to the motor torque constant, gear ratio, and current measured by the exoskeleton motors. Positive Power (+ P) was calculated using (3) and (4), where θ represents the angle of rotation of the motor. Similarly, peak positive power corresponds to the maximum of the positive power observed during the recording period. Average torque and average positive power were calculated by applying the average value formulas to both signals, shown in (5) and (6), where t is time and t_max_ is the maximum time of the data recording. For a given iteration average torque, peak positive power, and average positive power have a corresponding normalized metabolic cost value to which they were linearly regressed. To analyze the metabolic cost individually and across subjects, we used a linear regression to compare the metabolic cost in percent change compared to no exoskeleton to the average exoskeleton torque, peak exoskeleton power, and average positive exoskeleton power. Within these comparisons, the slope of the linear regression, Pearson’s correlation coefficient (r), and the p-value indicate trends between the metabolic cost and each presented metric.

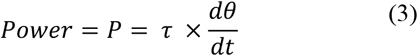

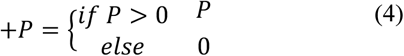

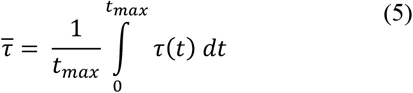

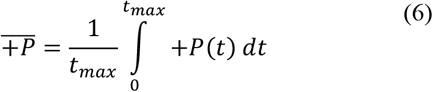

## III. Results

### A. Optimized Metabolic Cost vs. Subject-Task Dependency

The personalized control approach significantly (p<0.05) outperformed the no exoskeleton condition, with the only exceptions being the task agnostic and task dependent conditions during level ground walking at 1.5 m/s (Fig. 5). On average across all tasks, the personalized controller condition reduced metabolic cost by 18.3% which was significantly better (p<0.05) than both the task dependent (8.6% reduction) and task agnostic (8.4% reduction) conditions. For the individual tasks, all personalized control parametrizations reduced metabolic cost significantly (p<0.05) as compared to the task agnostic condition. Personalized control reduced metabolic cost significantly (p<0.05) as compared to the task dependent condition during level ground walking at 1.1 m/s (10.3% reduction), but the difference was not significant during level ground at 1.5 m/s (3.95% reduction) and the inclined walking (11.62% reduction) tasks. On average and in each individual walking task, there were no significant (p>0.05) differences between using the task agnostic or task dependent methods. Average metabolic landscapes for each tasks and across all tasks are shown in Fig. 6, with each subject’s best parameters and the average best parameters highlighted with colored triangles and stars, respectively.

**Fig. 5:**
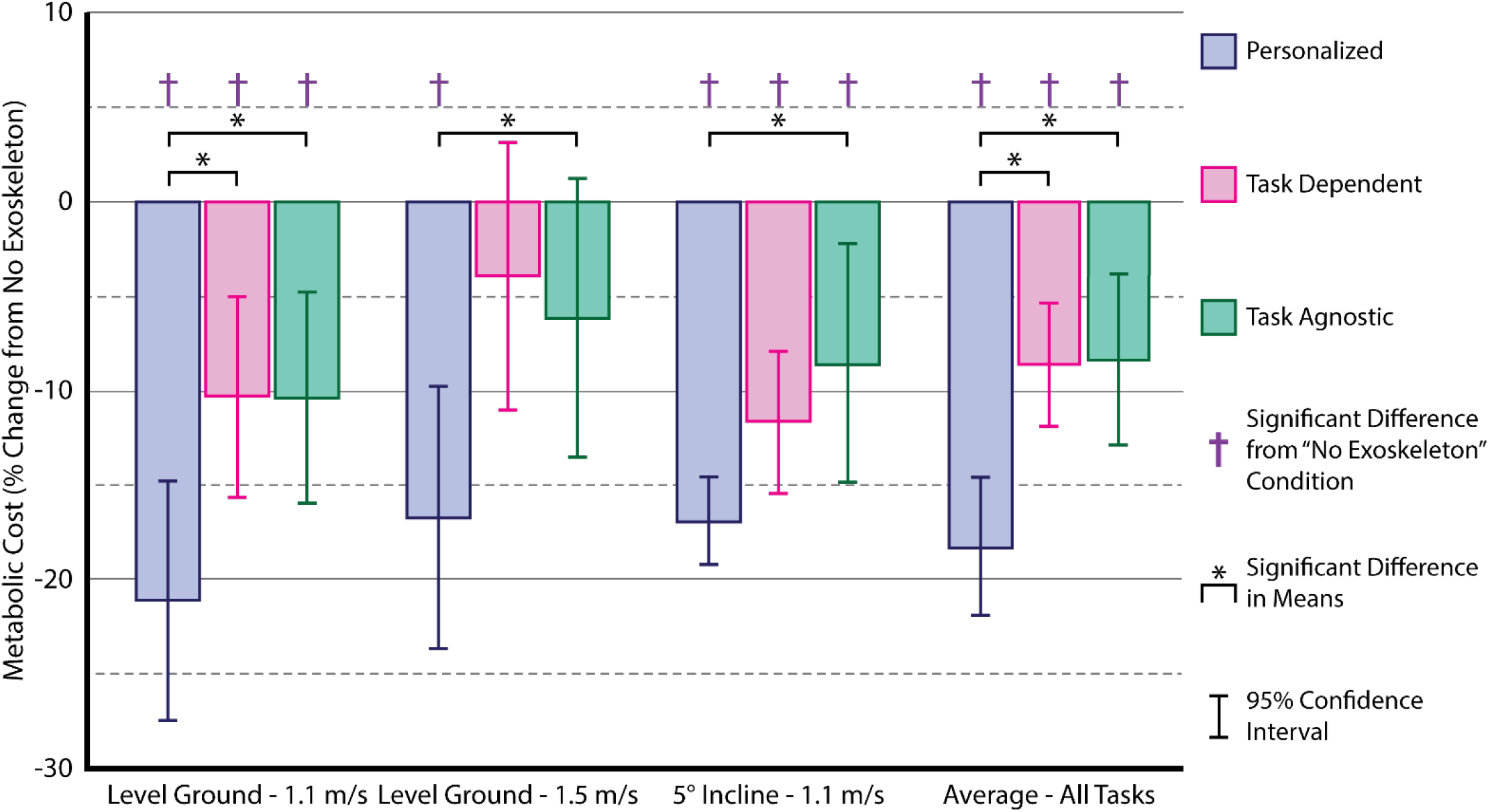
Comparisons between the subject-average metabolic benefit of the exoskeleton controller with respect to percent change from the subjects’ baseline metabolic cost when walking without the exoskeleton. Results are shown averaged across all ambulation modes as well as for each individual mode. Each bar represents 9 subjects’ metabolic cost at control parameters as determined by one of three methods. (i) Personalized control; (ii) Task Dependent control; and (iii) Task Agnostic control. A one-way ANOVA with a post hoc Bonferroni correction was used to determine the significant differences between metabolic cost within each ambulation mode, as well as the significant differences between each metabolic cost and the subjects’ no exoskeleton metabolic value. Notably, across all tasks and on average there is no significant difference between (ii) Task Dependent control and (iii) Task Agnostic control.

**Fig. 6:**
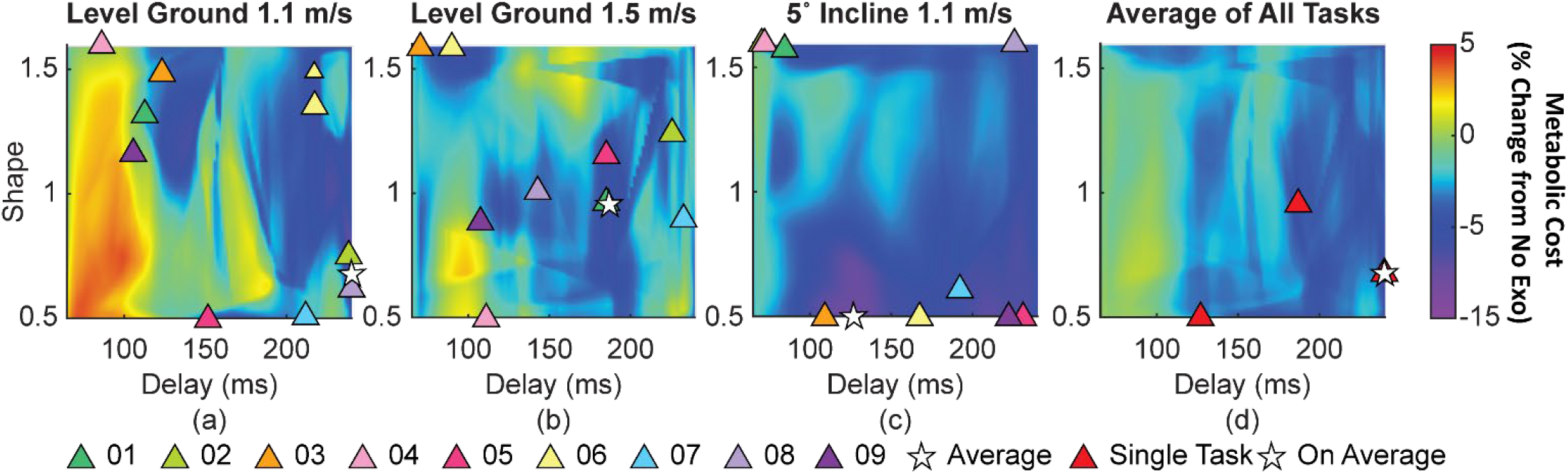
Average metabolic landscapes for **(a)** level ground walking at 1.1 m/s, **(b)** level ground walking at 1.5 m/s, **(c)** and 5° inclined walking at 1.1 m/s. The X and Y axes show the different exoskeleton parameters, and the color shows metabolic cost relative to the subjects’ baseline, no exoskeleton metabolic cost. Each surface seen here is the average of all nine subject’s individual metabolic landscape for the specified walking condition. Each triangle represents the parameters that correspond to the minimum metabolic cost of each subject’s individual metabolic landscape. The white star on each landscape represents the parameters that correspond to the minimum metabolic cost of the average surface across all subjects. **(d)** Metabolic landscape averaged over all three walking conditions. This surface represents the results from all 27 metabolic cost surfaces (9 subjects, 3 walking conditions). The red triangles represent the parameters that correspond to the minimum metabolic cost of each walking condition, while the white star represents the parameters that correspond to the lowest metabolic cost on the average surface of all walking conditions.

### B. Preference

Trends in the preference surfaces (Fig. 7) showed relative matches to those of the metabolic surfaces (Fig. 6) with a Pearson’s correlation coefficient of r = 0.69, indicating a relatively strong positive correlation. Across all ambulatory tasks, subjects had major bias towards quadrants I and IV, representing longer delays, with reasonably high uniformity. However, subjects preferred smaller shape values, such as quadrant IV over quadrant I, about 69% of the time at the slower walking speed, and during the higher walking speed trial subjects tended to prefer the higher shape quadrant around 57% of the time. For both level ground trials, subjects showed a strong bias against control parameters that have both a low delay and low shape, such as in quadrant III. Across all tasks, trends showed that users mostly preferred controller parameters within Quadrant IV, representing longer delay values and smaller shape values, between 59% and 74% of the time compared to other quadrants.

**Fig. 7:**
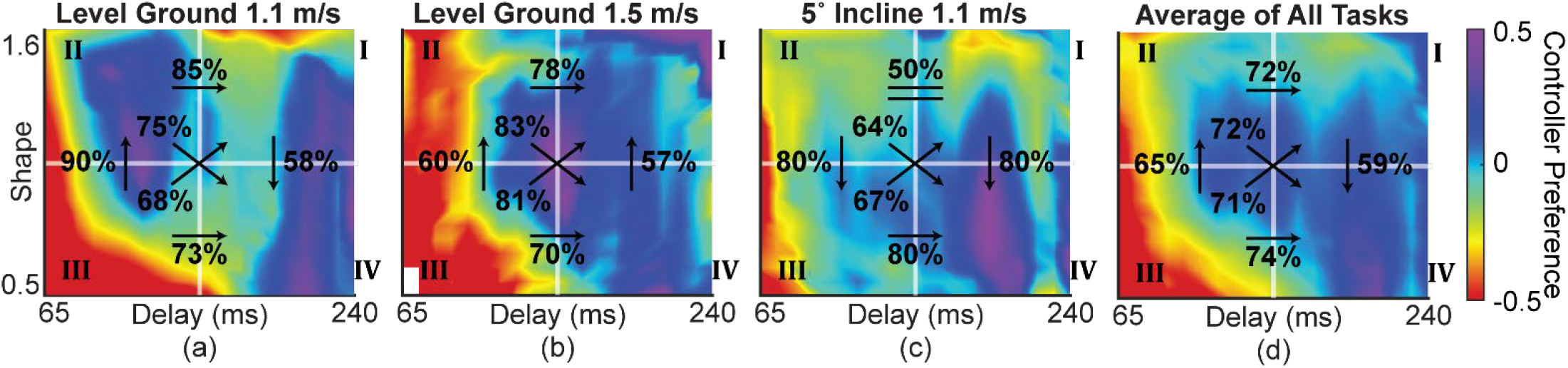
Preference landscapes for **(a)** level ground walking at 1.5 m/s**, (b)** level ground walking at 1.1 m/s, **(c)** inclined walking at 1.1 m/s, and **(d)** the average of all tasks using the pairwise comparison queried from subjects during the HILO experiment. Higher values (purple) are preferred over lower values (red). We separated the parameter space into four quadrants to analyze the frequency of which subjects preferred certain quadrants over others. The black arrows superimposed over the surfaces denote the quadrant analysis, where the direction of the arrow indicates quadrant preference, and the number adjacent to the arrow gives the percentage that quadrant was preferred over other quadrants.

### C. Exoskeleton Power & Torque

We used linear regression to observe the relationship between metabolic cost and exoskeleton assistance (Fig. 8). The linear regressor for metabolic cost and average exoskeleton torque had a slope of -1.379 percent-change per Nm/Kg (p<0.05), a Pearson’s correlation coefficient of r = -0.2007, and a statistically significant p value (p < 0.05). The linear regressor for metabolic cost and peak exoskeleton power had a slope of 0.0067 percent-change per W/Kg, a Pearson’s correlation coefficient of r = -0.0634, and a statistically insignificant p value (p > 0.05). Lastly, the linear regressor for metabolic cost and average exoskeleton power had a slope of 0.2985 percent-change per W/Kg, a Pearson’s correlation coefficient of r = -0.1653, and a statistically significant p-value (p < 0.05).

**Fig. 8:**
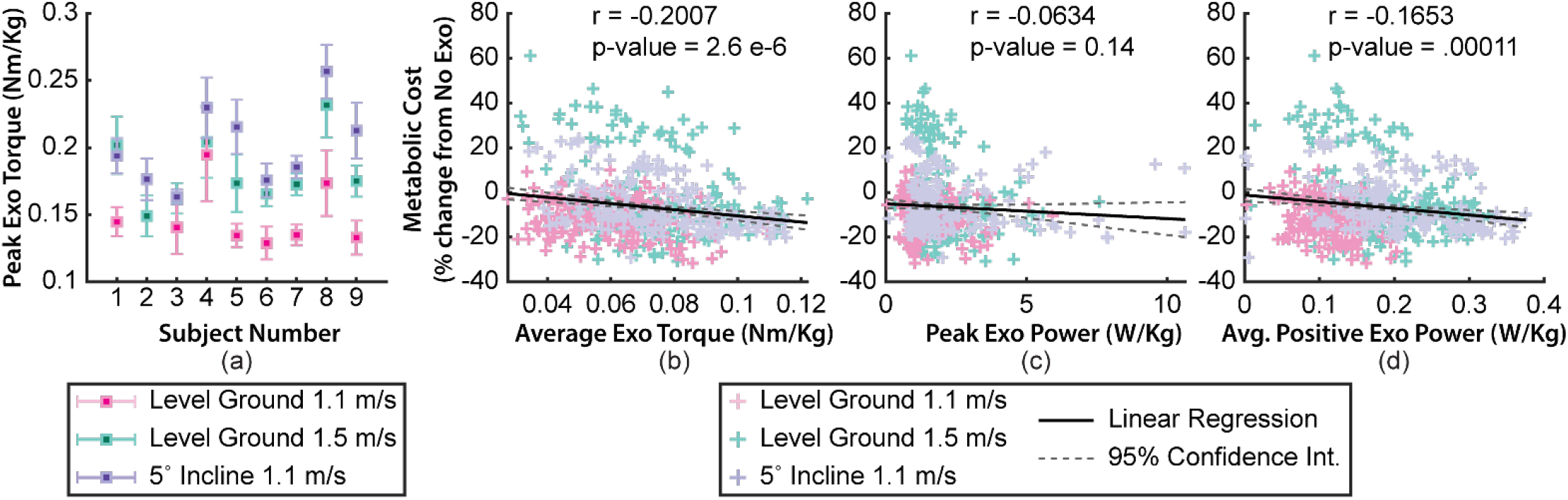
**(a)** Multi-modal peak exoskeleton torque for each subject. The goal was to keep the peak torque relatively constant across all iterations within each ambulation mode. Note: As subject two’s exoskeleton data was corrupted for their slower level ground mode, that data was not included in this analysis. Across 9 subjects with 21 iterations of differing control parameters and 3 ambulation modes, and withholding subject two’s missing data, there are 540 data points to relate exoskeleton torque and power data to subject metabolic cost. This data was used to regress metabolic cost versus **(b)** average torque provided by the exoskeleton, **(c)** peak power provided by the exoskeleton, and **(d)** average positive power provided by the exoskeleton. As expected, metabolic cost reduces as exoskeleton torque and power increase, however the p-values indicate that this relation is significant only for average exoskeleton torque and average positive exoskeleton power.

## IV. Discussion

This study investigated the effect of personalized exoskeleton assistance on the metabolic cost of 9 users across three cyclic tasks. Overall, our personalized control approach provided significant (p < 0.05) metabolic reductions across all three tested ambulation tasks when compared to both task dependent and task agnostic control approaches (Fig. 5), with the greatest metabolic reduction across all subjects and tasks accomplished during level ground walking at 1.1 m/s in which the subject average metabolic reduction was 21.1% when compared to the no exoskeleton condition. Our personalized control approach provided an average reduction in metabolic cost of 18.3%, whereas the task dependent approach provided an average reduction of 8.6%, thus verifying our primary hypothesis **I**. These results were expected based on previous literature that investigated similar personalized exoskeleton assistance via HILO when compared to generalized control parameters such as an end-to-end, deep learning driven biological torque-based controller [8], [9], [27], [30], [31], [32]. For example, the generalizable control approach presented in Molinaro *et al.* [16] on the same hip exoskeleton used in this study, provided a reduction in user metabolic cost of 5.4% and 10.3% for level ground and 5° inclined walking relative to the no exoskeleton condition, respectively. While this approach represents a novel method of exoskeleton control that can adapt to users in real-time, an essential requirement for translating exoskeletons out of the lab, the metabolic benefits of our personalized approach largely outperformed the purely generalizable controller across similar cyclic tasks, indicating the added value of personalizing exoskeleton control.

Across all modes, there was no statistically significant metabolic difference between using non-personalized task-dependent and task-agnostic controller settings (Fig. 5), thus rejecting hypothesis **II**. Overall, this indicates that for this type of biological moment-based hip exoskeleton controller in the absence of subject specific data, a controller setting that is task agnostic would be as robust as a controller that is task dependent across steady state, cyclic tasks. This is likely because in this controller paradigm there are areas of the landscape (specifically near Quadrant IV) that the user’s metabolic cost has low sensitivity to small changes in the controller parameters (Fig. 7), indicating that using parameters from a multi-task average metabolic landscape does not have a drastic metabolic penalty as compared to using single-task metabolic surfaces. When comparing the average metabolic and preference landscapes across both tasks and subjects, areas such as Quadrant IV have considerable overlap, with a relatively strong positive correlation (r = 0.69), indicating that as metabolic cost reductions increase, user preference similarly increases. For example, the most preferred set of parameters on average were found in Quadrant IV (shape = 0.5 and delay ≈ 212.72 ms) corresponding to a metabolic cost reduction of 6.23% on the average metabolic surface, suggesting that further HILO studies could improve personalized control methods by optimizing for user preference as a proxy for metabolic cost.

The controller parametrization used in this study was based on previously optimized spline-based hip exoskeleton control profiles observed during level ground walking and ramp ascent [8], [9], with our parametrization representing a profile that is flatter at times where the biological torque is near zero and sharper at peak biological torques. Offline optimizations based on this optimal torque profile showed that the addition of a shaping parameter, λ in (1), and a delay term to our parametrization enables our controller to further transform our estimated biological torque profile to a nearly identical profile to that of [9], with an R^2^ of 0.92 for level ground walking and R^2^ of 0.96 for ramp ascent. When tested on three expert-level users using a high-torque (∼200 Nm) tethered exoskeleton with only hip assistance, this previously validated optimal torque profile provided an average metabolic reduction of 6.72% relative to a no exoskeleton condition [9]. Our approach consisting of personalized exoskeleton assistance when deployed on a fully autonomous hip exoskeleton capable of providing 18 Nm of hip flexion and extension assistance, and across three cyclic tasks and 9 subjects with only 15 minutes of training, provided an average 18.3% reduction in user metabolic cost, representing an 11.6% difference in reductions when compared to the high-torque exoskeleton used in [9]. This highlights the importance of personalizing exoskeleton assistance on autonomous devices to maximize the benefits of assistive technology when deployed in the real world.

In our controller parametrization, we observed that the shaping term had a larger effect on the average power provided by the exoskeleton than the scaling and delay terms. This can be seen in Fig. 3, where lower shape values flatten the peak torques of the control profile, making the integration from (6) larger. Increasing positive power provided by an exoskeleton has been shown to improve metabolic cost reductions in users [22], thus optimizing the shape of the provided torque is highly important for personalized exoskeleton control. Likewise, when looking at the correlation between subject metabolic cost and average exoskeleton torque or positive power (Fig. 8), as exoskeleton average torque and power increases, metabolic cost significantly decreases, partially confirming our hypothesis **III**. It is interesting to note that peak exoskeleton power did not significantly correlate with metabolic cost, which may be due to the subjects trending towards higher average power as opposed to higher peak power delivered by the exoskeleton, i.e., as the shape term is increased there are higher peaks in the exoskeleton control profile, however the overall area under the profile’s curve is reduced. Similarly, at the slower ambulation tasks, such as level ground and incline walking at 1.1 m/s, the average best parameters lie in a place where average positive exoskeleton power delivered to the subject is very high. This benefit can be seen in the most physically demanding mode of the experiment at 5°-degree incline walking, where most metabolic reductions occurred at extremely low shape values during walking. Additional evidence is further reflected in user preference, where the level ground and inclined 1.1 m/s preference surfaces show quadrant IV having the strongest preference of all quadrants.

There are several limitations to this approach. First, human subjects’ experiments centered around metabolic cost are time-consuming and difficult to collect. Due to the strenuous nature of our experimental protocol, we limited metabolic trials to two-minute iterations, as opposed to the standard six-minute trials to ensure subjects were able to complete the entirety of the protocol, which may have reduced the accuracy of our metabolic landscapes. Secondly, our experiment focused only on steady-state cyclic activities, such as level-ground and inclined walking, which represents a small subset of the dynamic tasks and activities that humans perform throughout the day. While our mid-level control approach significantly reduced user metabolic cost during steady-state cyclic activities, it is difficult to know whether these results change for different or non-steady state ambulation modes such as running, sit-to-stands, stairs, or lifting weights without further investigation. In particular, the insensitivity of the metabolic cost landscape of the task agnostic controller may have more sensitivity outside of the tested cyclic locomotion tasks and requires further studies on the topic. Lastly, our exoskeleton was limited by the device hardware to provide a maximum of 18 Nm of assistive torque at the hip joint. While studies focused on high powered tethered exoskeletons have shown increased metabolic benefits for users, these are inherently difficult to translate outside of the lab due to their size and weight. Our device was autonomous and portable, providing relatively close metabolic reductions to these tethered devices while allowing for a more lightweight and autonomous device.

## V. Conclusion

This study used personalized exoskeleton control parameters derived from human-in-the-loop optimization to map the relationship between biological joint moment and optimal exoskeleton assistance. Overall, we found that our personalized control approach reduced the metabolic cost of walking by an average of 18.3% when compared to walking without an exoskeleton. Our results indicate that our personalized control approach can significantly reduce user metabolic cost across all three tested ambulation modes, as well as when compared to non-personalized task-dependent and task-agnostic control approaches. Furthermore, our results showed that there was no statistical significance between a task-dependent and task-agnostic control approach, suggesting that parameters derived from a multi-task metabolic HILO can potentially be used in a task-agnostic or generalizable control approach with minimal penalty to user metabolic cost. Our results further show that user preference has a relatively strong correlation with metabolic cost during steady state tasks, indicating the potential for future HILO studies to use user preference as a proxy for metabolic cost, thus avoiding time consuming and tiring optimization experiments. Furthermore, this approach can expand on end-to-end exoskeleton controllers such as a real-time biological joint moment estimator, to help improve and fine tune exoskeleton assistance for each user, representing a critical step to translating these devices to the real world.

